# Harnessing DNA Foundation Models for Cross-Species Transcription Factor Binding Site Prediction in Plant Genomes

**DOI:** 10.1101/2025.07.14.664780

**Authors:** Maryam Haghani, Krishna Vamsi Dhulipalla, Song Li

## Abstract

Accurate prediction of transcription factor binding sites (TFBSs) is crucial for understanding gene regulation. While experimental methods such as ChIP-seq and DAP-seq are informative, they are labor-intensive and species-specific. Recent advancements in large-scale pretrained DNA foundation models have shown promise in overcoming these limitations. This study evaluates the performance of three such models—DNABERT-2, AgroNT, and HyenaDNA—in predicting TFBSs in plants. Using DAP-seq data from *Arabidopsis thaliana* and *Sisymbrium irio*, we benchmark their accuracy against specialized approaches, including a motif-based method and two deep learning models, DeepBind and BERT-TFBS. Our results demonstrate that foundation models, particularly HyenaDNA, offer superior predictive accuracy and computational efficiency, highlighting their potential for scalable, genome-wide TFBS prediction in plants.

## 1 Introduction

Transcription factors (TFs) are key regulatory proteins that control gene expression by binding to specific DNA sequences called transcription factor binding sites (TFBSs). Accurate identification of these binding sites is crucial for deciphering gene regulatory networks and understanding biological processes. In plants, experimental methods such as Chromatin immunoprecipitation sequencing (ChIP-seq) [1] and DNA Affinity Purification sequencing (DAP-seq) [2] are commonly used to map TFBSs. While powerful, these assays are labor-intensive, costly, often limited to a few species, and lack the scalability needed for comprehensive, genome-wide analyses

The advent of machine learning and deep learning approaches has revolutionized TFBS prediction in animal and human systems by learning sequence features directly from high-throughput binding data [3]. However, their application in plant genomics has lagged behind, despite the availability of extensive DAP-seq and ChIP-seq data. A true paradigm shift has emerged with the rise of foundation models—large-scale pretrained models originally developed in natural language processing and now adapted to DNA sequences. These foundation models, including DNABERT-2, a transformer-based model pretrained on genomes from 135 species, demonstrating superior performance in various DNA sequence classification tasks [4]; the Agronomic Nucleotide Transformer (AgroNT) [5], a RoBERTa-style encoder pretrained on genomes from 48 plant species with state-of-the-art performance in regulatory feature prediction across plant genomes; and HyenaDNA [6], a decoder-only model utilizing Hyena operators for long-range genomic sequence modeling at single nucleotide resolution, offering efficient training and inference, have revolutionized genomics by capturing rich, complex representations of DNA sequences that generalize across species and tasks.

However, despite extensive datasets in plants, no prior study has systematically benchmarked multiple large pretrained DNA foundation models on predicting plant transcription factor binding site data from DAP-seq experiments, or evaluated their ability to generalize across species.

In this study, we leverage the power of these pretrained DNA foundation models by fine-tuning them on DAP-seq data from *Arabidopsis thaliana* (*A. thaliana*) to capture complex sequence patterns. We further evaluate model generalization through cross-species testing on the closely related *Sisymbrium irio* (*S. irio*). We selected these two species because they both belong to the Brassicaceae family, possess relatively small genomes, and exhibit highly similar—but not identical—transcription factor binding site landscapes for orthologous genes. Specifically, we focus on the ABF family of transcription factors, key mediators of abscisic acid (ABA)-regulated gene expression in plants [7]. ABA is a central plant hormone involved in mediating various stress responses. Understanding how ABFs bind across the genome can help identify more effective targets for enhancing plant tolerance to abiotic stress conditions, with the potential to improve crop yields during periods of environmental adversity such as drought. Due to the biological significance of ABF genes, they were the first among thousands of plant transcription factors for which cross-species binding sites were identified using DAP-seq experiments [7]. This provides a rich, biologically validated dataset for training and evaluating the predictive power of our foundation models. By appending a lightweight classification head and training end-to-end, our fine-tuned models learn nuanced sequence dependencies far beyond the reach of traditional TFBS predictors. We use a unified training and evaluation framework based exclusively on DNA sequences and systematically benchmark its performance and computational efficiency against established TFBS prediction approaches. Specifically, we compare with (1) a motif-based method utilizing MEME [8] and FIMO [9], and two deep learning models: (2) DeepBind [10], a convolutional neural network (CNN)–based predictor of DNA/RNA binding sites, and (3) BERT-TFBS [11], which integrates DNABERT-2 with CNN modules and a convolutional block attention module to capture both local and global sequence dependencies for TFBS prediction. Our results demonstrate that fine-tuned foundation models substantially outperform specialized predictors (motif-based, DeepBind and BERT-TFBS), and that HyenaDNA in particular achieves comparable predictive accuracy while reducing training time by over an order of magnitude. Finally, we conclude by discussing the limitations of our current methodology and outlining promising directions for future work.

## 2 Methods

### 2.1 Benchmark Datasets

We compiled DAP-seq binding site data for the ABA-Responsive Element Binding Factor (AREB/ ABF) family (AREB/ABF 1–4) from two public resources: `Malley2016` (Plant Cistrome Database)[2], which reports binding regions for multiple transcription factors in *Arabidopsis thaliana* (*A. thaliana*), including AREB/ABF2; and `Sun2022` [7], which characterizes AREB/ABF1–4 binding sites across four Brassicaceae species, most notably *A. thaliana* and *Sisymbrium irio* (*S. irio*). We focused on *A. thaliana*, which is a model organism in plant biology and human disease research [12]. Using these data, we defined three evaluation protocols:

1. **Cross-chromosome** (`Sun2022`): Leave-one-chromosome-out on *A. thaliana* AREB/ABFs across chromosomes 1–5 (14,374 unique samples in total). For each experiment, one chromosome serves as the test set and the remaining four as train/validation (see Supplementary Table S1). Because `Sun2022` sequences varied in length, we standardized them to maximum length of 265bp by padding shorter sequences.
2. **Cross-dataset** (`Malley2016` - `Sun2022`): Models were trained on AREB/ABF2 binding sites from `Malley2016` (*A. thaliana*) and evaluated on the corresponding AREB/ABF2 sites in `Sun2022`. Since `Malley2016` includes only AREB/ABF2, we extracted the same TF’s sites from `Sun2022`, which—being released later—provides an independent test set. After removing duplicate sequences across datasets, 14,294 unique samples remained (see Supplementary Table S2). All sequences were then standardized to 201bp (the `Malley2016` length) by trimming longer and padding shorter `Sun2022` sequences.
3. **Cross-species** (`Sun2022`): Train on *A. thaliana* and test on *S. irio* (and vice versa) for AREB/ABF 1–4. *A. thaliana* contributes 14,374 binding site samples; *S. irio* contributes 10,558; total = 24,932 across both species (see Supplementary Table S3). As in the cross-chromosome protocol, `Sun2022` sequences varied in length, we standardized them to the maximum observed length of 265bp by padding any shorter sequences.

For each evaluation protocol, we generated an equal number of negative samples for both the training and test sets to maintain balanced class proportions. Negative samples were generated by random permutation of nucleotides within each positive sequence, using a dinucleotide-preserving shuffling procedure that maintains both sequence length and dinucleotide composition. This approach disrupts functional motifs while retaining background compositional biases and is widely employed as a control strategy in TFBS prediction [10, 13, 14] (see Supplementary Section S1.2).

### 2.2 Model Architecture

To predict TFBS, we attach a two-neuron classification head directly atop the final token representations of each pre-trained DNA foundation model and fine-tune all model parameters—including the new head—on our labeled TFBS examples (Figure 1). These foundation models were initially trained via self-supervised learning on large-scale genomic corpora. We applied this approach to three different models:

**Figure 1.**
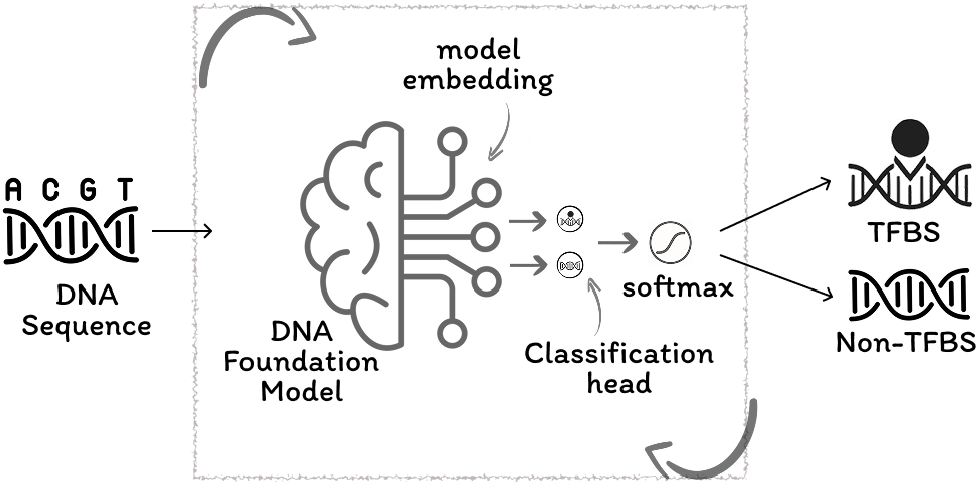
Overview of the fine-tuning framework for transcription factor binding sites (TFBS) prediction using a DNA foundation model: raw DNA sequences are input into a pre-trained DNA foundation model, which encodes them into high-dimensional embeddings. These embeddings are passed through a classification head that predicts whether the sequence corresponds to a TFBS or a non-TFBS. The loop arrows indicate that both the foundation model and the classification head are jointly updating the entire model end-to-end for the TFBS prediction task.

#### AgroNT

Building upon Nucleotide Transformer [15], Agronomic Nucleotide Transformer (AgroNT) is a 1B-parameter RoBERTa-style [16] encoder pre-trained on genomes from 48 plant species. It uses a 6-mer tokenizer and 40 transformer layers that generate 1,500-dimensional embeddings per token. The model was evaluated on a range of prediction tasks, including regulatory feature detection and gene expression, and demonstrated state-of-the-art performance [5]

#### DNABERT-2

A modern foundation model, with 117 million parameters, designed for modeling genomic sequences in multiple species. It uses Byte Pair Encoding (BPE) [17] tokenization over multi-species genomic sequences (32.5B bases) of 135 species and a 12 layer BERT encoder producing 768 dimensional embeddings [4]. Following sequence encoding, we apply mean pooling over token embeddings and feed the result into our two-class output head.

#### HyenaDNA

A decoder-only model that uses Hyena operators [18]—implicit long-range convolutional filters—to process up to 1M nucleotide tokens at single-base resolution. Its architecture forgoes k-mer tokenization in favor of single-nucleotide tokens and is trained via next-token prediction [6]. Its ability to model long-range interactions while maintaining single-nucleotide precision enables it to generalize effectively across tasks and species, establishing HyenaDNA as a robust and efficient foundation model for genomics [6].

### 2.3 Baseline Models

We employed several baselines to benchmark the performance and efficiency of our fine-tuning approach. In particular, we considered two recent deep learning architectures specifically developed for the binding-site prediction task:

**DeepBind** [10] uses a four-stage CNN to predict DNA- and RNA-binding specificity without manual feature engineering. First, raw nucleotide sequences are transformed into one-hot encoded matrices, which convolutional layers then scan using learned motif detectors to extract informative patterns. Second, these feature maps are passed through a rectification layer—applying an absolute value operation—to emphasize signal, then through a pooling layer that both enhances feature robustness and reduces overfitting. Finally, the condensed features feed into a fully connected layer that calculates a binding probability score. For our classification, we apply a softmax to this score to yield TFBS versus non-TFBS probabilities.

**BERT-TFBS** [11] integrates three modules: (1) a pretrained DNABERT-2 [4] encoder to capture long-range dependencies, (2) a CNN stack to extract high-order local features from the sequence, and (3) a Convolutional Block Attention Module (CBAM) [19] to enhance local features by the spatial and channel attention mechanisms on the convolutional outputs. The output module integrates all the sequence features from the three modules and passes them through a multi-layer perceptron output head for binary TFBS classification.

In addition to the deep learning baselines, we implemented a motif-based method using the MEME Suite [20]. Training set binding sequences were provided to the MEME motif discovery tool [8], which identified five de novo motifs represented as Position Weight Matrices (PWMs). Each PWM encodes nucleotide probabilities at individual motif positions. We then used FIMO [9], also part of the MEME Suite [20], to score all test sequences based solely on their similarity to the discovered motifs. This motif-centric baseline enables direct comparison with models capable of learning more complex sequence patterns.

## 3 Results

We evaluated our fine-tuned models (Section 2.2) alongside baseline models (Section 2.3) across three distinct experimental scenarios (Section 2.1) to benchmark TFBS prediction performance.

For all models, hyperparameters were set to each architecture’s default values. Training employed binary cross-entropy loss optimized via AdamW. Early stopping was triggered after 10 epochs without validation loss improvement, with the checkpoint exhibiting the lowest validation loss retained for testing. Performance was comprehensively measured using five metrics: accuracy (ACC), F1 score, Matthews correlation coefficient (MCC), area under the receiver operating characteristic curve (ROC-AUC), and area under the precision-recall curve (PR-AUC). This fine-tuning and evaluation pipeline facilitates a direct comparison between specialized TFBS predictors and pretrained genomic models. Implementation details are described in Supplementary Section S2. The following presents the results for each of the three evaluation scenarios.

### 3.1 Cross-Chromosome Evaluation on *A. thaliana* AREB/ABF1-4 Binding Sites

To assess each model’s ability to generalize to unseen genomic contexts, we evaluated them using a leave-one-chromosome-out scheme on AREB/ABF1-4 transcription factor binding sites of *A. thaliana* species from `Sun2022` dataset. In this protocol, each of the five *A. thaliana* chromosomes is held out in turn for testing, while the remaining four serve for training and validation. Table 1 summarizes the macro-mean performance metrics, as well as running times, aggregated over all 25 runs. The distribution of these metrics and runtimes for each held-out chromosome is shown in Supplementary Figure S1 (see Supplementary Table S6 for the per-chromosome averages across the five seeds used to train independent models).

**Table 1:** Macro-mean performance metrics for AREB/ABF transcription factor binding site regions in *A. thaliana*, evaluated under a **leave-one-chromosome-out** protocol. Each of the five chromosomes is used once as test set, with models trained using five different random seeds per setting. Reported metrics are averaged across all 25 runs (5 chromosomes × 5 seeds), with the best score in each column shown in bold.

| Model | ACC | F1 | MCC | ROC-AUC | PR-AUC | Train Time(s) | Test Time(s) |
| --- | --- | --- | --- | --- | --- | --- | --- |
| Motif-based | 0.738 | 0.687 | 0.505 | 0.750 | 0.734 | 3505.0 | 6.9 |
| DeepBind | 0.880 | 0.880 | 0.762 | 0.948 | 0.949 | 969.5 | 4.5 |
| BERT-TFBS | 0.904 | 0.916 | 0.808 | 0.947 | 0.949 | 5,600 | 28.8 |
| DNABERT-2 | 0.914 | 0.915 | 0.831 | 0.970 | 0.964 | 5,527 | 28.2 |
| AgroNT | **0.961** | **0.961** | **0.924** | **0.991** | **0.992** | 33,669 | 194.9 |
| HyenaDNA | 0.940 | 0.939 | 0.880 | 0.981 | 0.982 | **261.4** | **4.0** |

Our evaluation shows a clear progression from motif-based to deep learning approaches. The motif baseline yielded the weakest results despite a nontrivial training cost, while DeepBind achieved moderate performance (ACC/F1 ∼0.88) with reasonable efficiency. Transformer-based models (BERT-TFBS, DNABERT-2) further improved accuracy but at considerable computational expense. AgroNT delivered the strongest overall performance, though at the expense of extremely long training and inference times. By contrast, HyenaDNA reached near–state-of-the-art accuracy while training ∼130× faster and inferring ∼50× faster than AgroNT. This remarkable combination of near–state-of-the-art accuracy and minimal computation makes HyenaDNA particularly well-suited for large-scale, genome-wide binding-site prediction.

### 3.1 Cross-Dataset Evaluation on *A. thaliana* AREB/ABF2 Binding Regions

To assess cross-dataset generalization, we trained on *A. thaliana* AREB/ABF2 peaks from the `Malley2016` dataset and evaluated on the corresponding *A. thaliana* AREB/ABF2 peaks in `Sun2022` (see Section2.1, cross-dataset (`Malley2016` - `Sun2022`)).

Figure 2 (a) presents the test performance distributions in the cross-dataset setting for six models trained with different random seeds; corresponding mean values are reported in Supplementary Table S7. Statistical analysis using ANOVA followed by Tukey’s HSD test indicates that models annotated with the same letter (shown above the plot) are not significantly different at the 0.05 significance level. While no significant differences in accuracy, F1, or MCC were observed among the neural models, all consistently outperformed the motif-based baseline, which exhibited significantly lower ROC-AUC and PR-AUC values. Wilcoxon tests confirmed HyenaDNA’s superior accuracy, F1, and MCC over all other methods (see Supplementary Section S3.2), highlighting its clear advantage in cross-dataset generalization. Figure 2 (b) compares training and inference times across models with different random seeds (see Supplementary Table S7). AgroNT was the slowest to train, followed by motif-based, DNABERT-2, and BERT-TFBS, while HyenaDNA trained nearly 90× faster than AgroNT. DeepBind also trained quickly but with lower accuracy (Figure 2 (a)). For inference, HyenaDNA matched DeepBind as the fastest, clearly outperforming other neural models. Overall, these results highlight that HyenaDNA offers the best balance of speed and accuracy, making it ideal for large-scale cross-dataset use.

**Figure 2.**
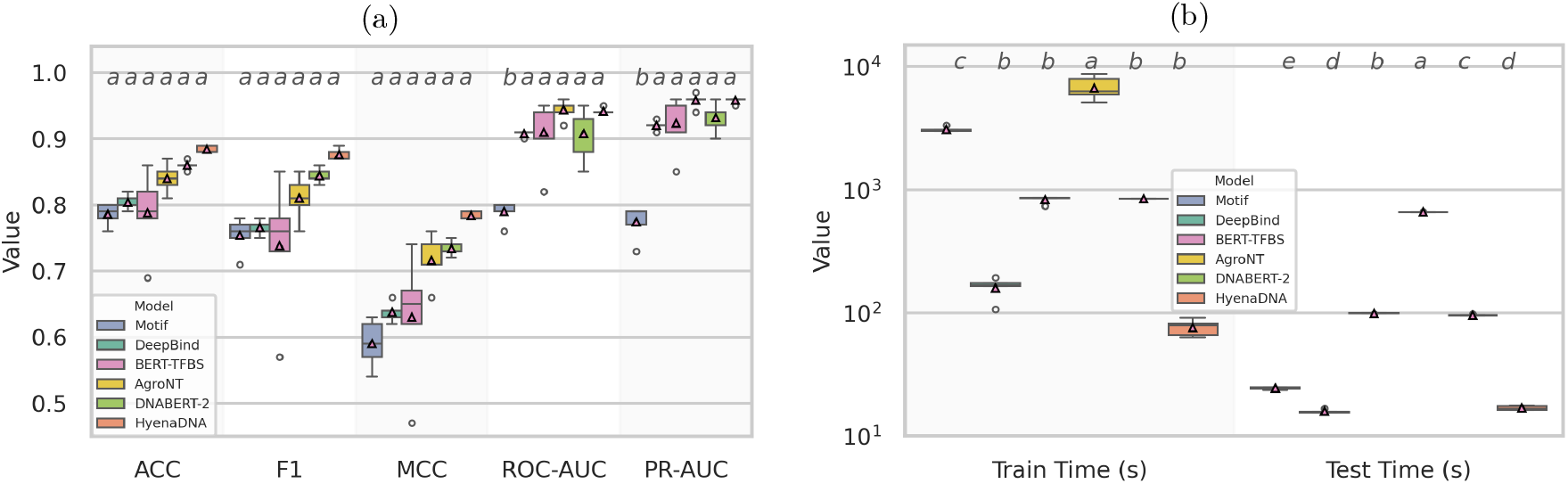
Distribution of **(a)** test-set performance metrics and **(b)** training and inference time (seconds). Models were trained with 5 random seeds on the `Malley2016` AREB/ABF2 dataset and evaluated on the `Sun2022` *A. thaliana* AREB/ABF2 test set in a **cross-dataset setting**. Letters above boxes indicate statistical groupings; models sharing a letter are not significantly different at the 0.05 level.

### 3.3 Cross-Species Transferability of AREB/ABF1-4 Binding Sites

To evaluate how well our models generalize beyond a single species, we assembled a cross-species dataset of AREB/ABF 1–4 DAP-seq peaks from two Brassicaceae: *A. thaliana* and its wild relative *S. irio* from `Sun2022`. We then carried out a leave-one-species-out evaluation—training once on *A. thaliana* and testing on *S. irio*, then vice versa. Figure 3 shows the distributions of test-set performance metrics and runtimes, with mean values reported in Supplementary Table S8 across five independently trained models per transfer direction.

**Figure 3.**
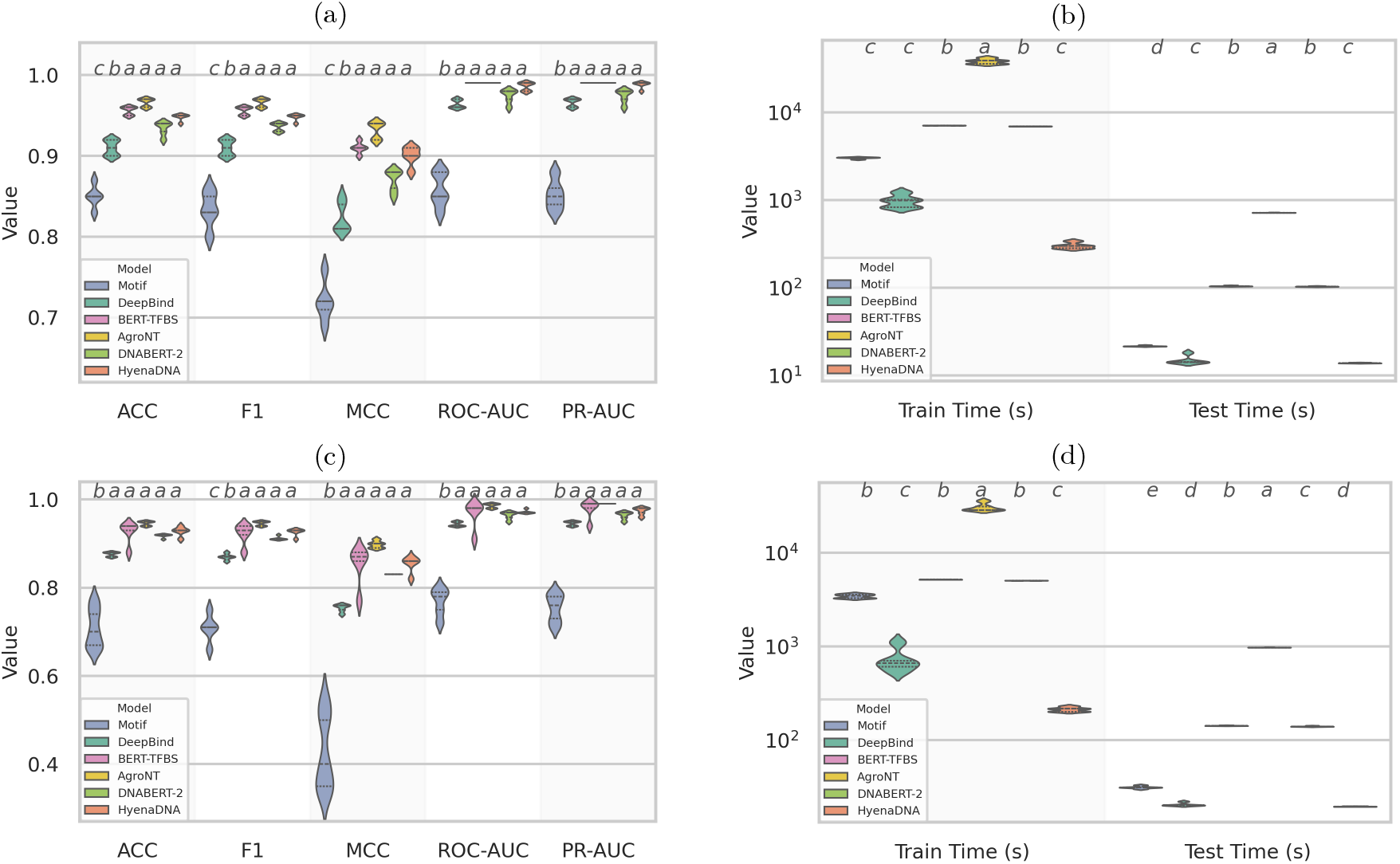
Distribution of test-set performance metrics as well as training and inference times (in seconds) for models in the **cross-species evaluation**. Each distribution is based on five independently trained models with different random seeds, evaluated on the corresponding test species. **(a–b)**: models trained on *A. thaliana* and tested on *S. irio*; **(c–d)**: models trained on *S. irio* and tested on *A. thaliana*. Letters above boxes indicate statistical groupings; models sharing the same letter are not significantly different at the 0.05 level.

Despite the phylogenetic distance between *A. thaliana* and *S. irio*, deep learning models generalized well across species, whereas the motif-based baseline showed limited predictive power (mean MCC: 0.720 trained on *A. thaliana*, 0.426 on *S. irio*) and required substantially more training time (>3,000s vs.302s/213s for HyenaDNA). Among neural models, ANOVA followed by Tukey’s HSD test confirmed that HyenaDNA achieved performance comparable to all others for ROC-AUC and PR-AUC, while matching DNABERT-2, BERT-TFBS, and AgroNT for accuracy, F1, and MCC.

AgroNT achieved slightly higher accuracy, F1, and MCC in both transfer directions, but at extreme cost (>38,000s and 30,000s training; with inference up to 977s). By contrast, HyenaDNA offered competitive accuracy at far lower runtime—over 100× faster than AgroNT and 20× faster than BERT-TFBS and DNABERT-2 in training, with similarly large gains in inference speed. These results highlight HyenaDNA’s strong balance of accuracy and efficiency in cross-species prediction.

This robust cross-species performance is underpinned by the high conservation of AREB/ABF binding motifs: Sun et al. (2022) showed that the DNA-binding preferences of AREB/ABF transcription factors are essentially conserved across Brassicaceae species, including *A. thaliana* and *S. irio*, and comparison of the position weight matrices of all AREB/ABFs revealed little difference in binding site preference. Also, swap-DAP assays revealed extensive overlap, indicating that TF–DNA contacts are conserved despite species divergence [7]. This high degree of motif and binding-site conservation provides a mechanistic justification for why a model trained on one species can accurately predict AREB/ABF binding regions in the other.

### 3.4 Ablation Study on HyenaDNA

In this section, we conduct an ablation analysis of HyenaDNA to better understand how each component of the model contributes to its overall performance. We selected this model for its state-of-the-art predictive accuracy and exceptional efficiency among all fine-tunable models. Supplementary Table S9 lists the trainable parameters in HyenaDNA under the three freezing regimes. When the backbone is fully frozen (`Backbone`), only the 258-parameter classification head (<0.1%) is updated. Allowing embeddings and the classification head to train (`Backbone.layers`) increases the parameters to <0.6% of all parameters in the full model (represented by `None` in Table 2).

**Table 2:** Performance metrics of HyenaDNA ablation experiments across different evaluation scenarios. Each sub-table displays the mean values of the performance metrics.

(a) Macro-mean performance for **cross chromosome evaluation** of AREB/ABF1-4 in *A. thaliana*
| Frozen Component | ACC | F1 | MCC | ROC-AUC | PR-AUC |
| --- | --- | --- | --- | --- | --- |
| backbone | 0.633 | 0.640 | 0.269 | 0.686 | 0.670 |
| backbone.layers | 0.864 | 0.868 | 0.733 | 0.942 | 0.946 |
| none | 0.940 | 0.940 | 0.880 | 0.981 | 0.982 |

(b) Mean performance for **cross-dataset evaluation** (AREB/ABF2; trained on Malley2016, tested on Sun2022)
| Frozen Component | ACC | F1 | MCC | ROC-AUC | PR-AUC |
| --- | --- | --- | --- | --- | --- |
| backbone | 0.606 | 0.632 | 0.216 | 0.650 | 0.632 |
| backbone.layers | 0.794 | 0.786 | 0.592 | 0.874 | 0.882 |
| none | 0.884 | 0.876 | 0.784 | 0.942 | 0.958 |

(c) Mean performance for **cross-species evaluation** for AREB/ABF1-4 for *A. thaliana* and *S. irio*.
| Split | Frozen Component | ACC | F1 | MCC | ROC-AUC | PR-AUC |
| --- | --- | --- | --- | --- | --- | --- |
| **train:***A. thaliana*<br>**test:***S. irio*. | backbone | 0.630 | 0.632 | 0.260 | 0.680 | 0.670 |
|  | backbone.layers | 0.878 | 0.884 | 0.760 | 0.952 | 0.954 |
|  | none | 0.948 | 0.948 | 0.900 | 0.986 | 0.988 |
| **train:***S. irio*.<br>**test:***A. thaliana* | backbone | 0.634 | 0.612 | 0.272 | 0.690 | 0.670 |
|  | backbone.layers | 0.868 | 0.870 | 0.742 | 0.942 | 0.944 |
|  | none | 0.928 | 0.926 | 0.854 | 0.972 | 0.974 |

Tables 2a–2c compare these regimes across cross-chromosome, cross-species and cross-dataset tasks. In every case, updating only the embeddings and classification head (`Backbone.layers`) delivers performance nearly identical to full fine-tuning while adjusting under 0.6% of parameters. These results demonstrate HyenaDNA’s parameter-efficient adaptability: by freezing over 99% of its weights, it maintains high accuracy and AUC, substantially reducing both computational load and memory usage with minimal impact on predictive power. Supplementary Figures S2-4 show the full distribution of metrics for each evaluation scenario.

## 4 Conclusion

This study demonstrates the superior performance of large-scale pre-trained DNA foundation models in predicting transcription factor binding sites (TFBSs) in plants. Our evaluations on *Arabidopsis thaliana* and *Sisymbrium irio* datasets reveal that these models, particularly HyenaDNA, outperform traditional architectures in both predictive accuracy and computational efficiency.

Looking ahead, our main downstream objective is to develop a model for transcription factor binding site (TFBS) prediction that can impute unperformed DAP-seq experiments by inferring peaks from promoter regions in species lacking experimental data. This will enable the expansion of our analysis to a broader range of plant species and allow assessment of model generalizability across diverse genomes, providing a scalable framework for cross-species prediction. At present, TFBS data for the same transcription factor (ABFs) are available for only a limited number of species within the Brassicaceae family. Upcoming multi-DAP-seq experiments [21] are expected to generate rich datasets that will enable further testing of the computational pipeline developed in this work. To improve predictive performance, we also plan to incorporate transcription factor–specific information, enabling the development of TF-aware models capable of transferring knowledge across distinct TFs. In parallel, the integration of interpretation methods will support the transfer of regulatory network knowledge from well-studied model plants to less-characterized species. Collectively, these advances will support scalable, genome-wide TFBS prediction and broaden our understanding of gene regulation across diverse plant genomes.

## 5 Code and Data availability

All scripts are available in our GitHub repository: https://github.com/Maryam-Haghani/TFBS The processed data are deposited in Zenodo: https://zenodo.org/records/17229680 (DOI: 10.5281/zenodo.17229680).

