## Supplementary File for "Harnessing DNA Foundation Models for Cross-Species Transcription Factor Binding Site Prediction in Plant Genomes"

#### S1 Dataset

##### S1.1 Statistics

Tables S1 - S2 summarize the dataset statistics for the one-chromosome-held-out, species-held-out, and cross-dataset experiments, respectively.

Table S1: Distribution of `Sun2022` AREB/ABF transcription factor binding site samples across different chromosomes for *Arabidopsis thaliana* (*A. thaliana*).

| Chromosome | # samples |
|---|---|
| Number of peak samples | 14,454 |
| Number of unique peak samples | 14,374 |
| Chromosome 1 | 3,787 |
| Chromosome 2 | 2,232 |
| Chromosome 3 | 2,807 |
| Chromosome 4 | 2,297 |
| Chromosome 5 | 3,251 |

Table S2: Dataset statistics for AREB/ABF2 binding sites in *A. thaliana* used in the cross-dataset evaluation. 735 duplicate records denote overlapping sequences between `Malley2016` and `Sun2022`; unique samples reflect the total after deduplication.

| Statistic | Malley2016 | `Sun2022` |
|---|---|---|
| Number of peak samples | 2,299 | 13,005 |
| Unique samples after deduplication | 2,299 | 11,995 |
| Mean sequence length | 201 | 201 |
| Standard deviation | 0 | 0 |
| Minimum length | 201 | 201 |
| 25% length | 201 | 201 |
| 50% length (median) | 201 | 201 |
| 75% length | 201 | 201 |
| Maximum length | 201 | 201 |

Table S3: Dataset statistics of `Sun2022` AREB/ABF transcription factor binding sites for *Arabidopsis thaliana* (*A. thaliana*) and *Sisymbrium irio* (*S. irio*) used in cross-species experiment.

| | *A. thaliana* | *S. irio* |
|---|---|---|
| Number of peak samples | 14,454 | 10,600 |
| Number of unique peak samples | 14,374 | 10,558 |
| Mean | 212.13 | 209.58 |
| Standard Deviation (std) | 5.33 | 3.02 |
| Minimum (min) | 208 | 169 |
| 25% | 210 | 208 |
| 50% (Median) | 210 | 209 |
| 75% | 211 | 209 |
| Maximum (max) | 265 | 241 |

##### S1.2 Negative set

We generated an equal number of negative samples to match the positive set by applying dinucleotide shuffling with uShuffle [1]. This approach has been a consistent and significant practice in the field of transcription factor binding site (TFBS) prediction since 2015, particularly with the rise of deep learning models [2, 3, 4]. Dinucleotide shuffling is particularly advantageous because it creates background sequences that control for lower-order sequence biases, such as GC content, by preserving the dinucleotide frequency of the positive (binding) sequences. As a result, predictive models are compelled to capture the true sequence motifs and patterns associated with transcription factor binding, rather than relying on simple compositional differences.

#### S2 Implementation Details

For the motif-based approach, positively labeled sequences from the training set were provided as input to MEME. We used discriminative mode using negatively labeled (unbound) sequences to identify motifs enriched in the positive set relative to the negative set. FIMO was then used on the test set with a significance threshold parameter set to 0.05, which defines the minimum match strength required for a subsequence to be classified as a putative binding site.

All deep learning models, whether classic architectures or pretrained DNA foundation models, were trained or fine-tuned according to the following unified protocol. This unified fine-tuning/training pipeline enables a direct comparison between specialized TFBS architectures and large-scale pretrained genomic models.

- **Model setup**
  - Classic models:
    - **DeepBind**: implemented via the MLSNET codebase [5].
    - **BERT-TFBS**: obtained from the original GitHub repository [6].
  - Pretrained DNA foundation models:
    - Removed any existing language-modeling head.
    - Appended a new, randomly initialized linear classification layer with two outputs (TFBS vs. non-TFBS).
- **Training Configuration**
  - **Loss function:** Binary cross-entropy

– **Optimizer:** AdamW [7].
– **Regularization:** Dropout applied to mitigate overfitting [8].
– **Batching:** All sequences were padded or truncated to the dataset's maximum input length.
– **Parameter updates:** All layers, including embeddings, were fine-tuned. The total number of trainable parameters for each model is listed in Supplementary Table S4.

Table S4: Number of trainable parameters for each model.

| **Model** | **# Parameters** |
|---|---|
| DeepBind | 2,194 |
| DNABERT-2 | 117,070,082 |
| BERT-TFBS | 142,319,622 |
| AgroNT | 985,099,603 |
| HyenaDNA (hyenadna-tiny-1k-seqlen) | 436,354 |

– **Hyperparameters:** Learning rates, batch sizes, and other hyperparameters were set to each model's default values (see Supplementary Table S5). When hyperparameter tuning was required, we employed stratified 5-fold cross-validation on the training set to maintain balanced class proportions across folds. In each iteration, one fold was used for validation, while the remaining four were used for model training.

Table S5: Default hyperparameter settings for each model. Epoch counts for fine-tuning were chosen empirically.

| Model | Kernel size | Epochs | Adj. LR | Train BS | Eval BS | LR | Weight Decay |
|---|---|---|---|---|---|---|---|
| DeepBind | 24 | 1,000* | No | 64 | – | $1 \times 10^{-4}$ | $1 \times 10^{-4}$ |
| BERT-TFBS | – | 15 | Yes (max $1.5 \times 10^{-5}$, min $2 \times 10^{-6}$) | 32 | – | $1.5 \times 10^{-5}$ | $1 \times 10^{-2}$ |
| DNABERT-2 | – | 15 | No | 32 | – | $1.5 \times 10^{-5}$ | $1 \times 10^{-2}$ |
| AgroNT | – | 60 | No | 8 | 64 | $1 \times 10^{-5}$ | $1 \times 10^{-1}$ |
| HyenaDNA | – | 60 | No | 32 | – | $1 \times 10^{-4}$ | $1 \times 10^{-1}$ |

*Although DeepBind's default # of epochs is 1,000, training never exceeded 100 epochs due to early stopping.

- **Checkpointing**
  – 5% of the training data was reserved as a hold-out validation set, with stratification applied to preserve balanced class distributions across the splits.
  – Validation loss was monitored each epoch, with early stopping after 10 epochs of no improvement.
  – The checkpoint yielding the lowest validation loss was retained for testing.

##### S2.1 Evaluation Metrics

Because TFBS prediction is a binary classification task, we evaluate each model using five complementary metrics—accuracy (ACC), F1 score, Matthews correlation coefficient (MCC), ROC-AUC and PR-AUC— to provide a comprehensive assessment of classifier performance:

- **Accuracy** [9]: fraction of correctly classified samples, including TFBSs and non-TFBSs, to all the tested samples.

$$\text{Accuracy} = \frac{TP + TN}{TP + FN + TN + FP}$$

where TP, FN, TN, and FP denote the number of true positives, false negatives, true negatives, and false positives, respectively.

- **F1 score**: harmonic mean of precision (Pr) and recall (Re)

$$F1 = 2 \times \frac{\mathrm{Pr} \times \mathrm{Re}}{\mathrm{Pr} + \mathrm{Re}}$$

- **Matthews correlation coefficient (MCC) [10]**: balanced correlation coefficient, robust under imbalance

$$\mathrm{MCC} = \frac{TP \times TN - FP \times FN}{\sqrt{(TP + FP)(TP + FN)(TN + FP)(TN + FN)}}$$

- **ROC-AUC** [11]: area under the receiver operating characteristic (ROC) curve, which plots

$$\mathrm{TPR} = \frac{TP}{TP + FN}, \quad \mathrm{FPR} = \frac{FP}{FP + TN}$$

over all thresholds. It is suitable for evaluating the performance of classifiers at different operating points.

- **PR-AUC** [12]: area under the precision–recall (PR) curve, trading off

$$\mathrm{Precision} = \frac{TP}{TP + FP}, \quad \mathrm{Recall} = \frac{TP}{TP + FN}$$

and more sensitive to class imbalance.

### S3 Results

#### S3.1 Cross-Chromosome Evaluation on *A. thaliana* AREB/ABF1-4 Binding Sites

Figure S1 illustrates the distributions of test performance and training cost under a leave-one-chromosome-out scheme. For each of the five A. thaliana chromosomes, one chromosome was reserved as the test set while models were trained on the remaining four. Distributions summarize results from five models initialized with different random seeds, reporting ACC, F1 score, MCC, ROC-AUC, PR-AUC, and total training time. AgroNT consistently achieves the highest absolute values for ACC, F1, MCC, ROC-AUC, and PR-AUC. Statistical analysis using ANOVA followed by Tukey's HSD test reveals that models sharing the same letter are not significantly different at the 0.05 significance level. This analysis shows that AgroNT, HyenaDNA, DNABERT-2, and BERT-TFBS perform comparably across all metrics and chromosome splits, often sharing the same significance group. DeepBind provides moderate improvements over the motif-based baseline, but it remains significantly weaker than transformer-based approaches, while motif-based methods are the least effective and highly variable. For computational efficiency, the contrast is stark: AgroNT is the most resource-intensive, requiring by far the longest training times, whereas HyenaDNA is exceptionally efficient, training almost 130 times faster and running inference nearly 50 times faster than AgroNT while still achieving competitive performance metrics. Overall, HyenaDNA offers the best balance between performance and computational cost.

Table S6 reports per-chromosome mean performance and training time for each model (motif-based, BERT-TFBS, DNABERT-2, DeepBind, HyenaDNA, AgroNT) under the leave-one-chromosome-out protocol. For each held-out chromosome, results are averaged over five models trained with different random seeds, highlighting variation in accuracy, correlation, AUC scores, and computational cost across distinct genomic contexts.

Table S6: Per-chromosome mean performance and timing for each model under **leave-one-chromosome-out** evaluation. Metrics include accuracy (ACC), F1 score, MCC, ROC-AUC, PR-AUC, total training time, and inference time. Each entry is the average over five models trained with different random seeds on the held-out chromosome. For each split, the top value in each column is highlighted in bold.

| Inference Chromosome | Model | ACC | F1 | MCC | ROC-AUC | PR-AUC | Train Time(s) | Test Time(s) |
|---|---|---|---|---|---|---|---|---|
| chromosome 1 | Motif-based | 0.780 | 0.764 | 0.570 | 0.804 | 0.796 | 2,800 | 9 |
| | DeepBind | 0.882 | 0.884 | 0.768 | 0.950 | 0.950 | 908 | 6 |
| | BERT-TFBS | 0.934 | 0.934 | 0.870 | 0.990 | 0.990 | 5,205 | 38 |
| | DNABERT-2 | 0.920 | 0.918 | 0.836 | 0.974 | 0.970 | 5,100 | 37 |
| | AgroNT | **0.968** | **0.968** | **0.936** | **0.992** | **0.994** | 29,169 | 256 |
| | HyenaDNA | 0.950 | 0.950 | 0.898 | 0.988 | 0.988 | **215** | **5** |
| chromosome 2 | Motif-based | 0.718 | 0.676 | 0.450 | 0.736 | 0.724 | 3,535 | 6 |
| | DeepBind | 0.866 | 0.864 | 0.728 | 0.934 | 0.938 | 335 | 4 |
| | BERT-TFBS | 0.914 | 0.908 | 0.832 | 0.972 | 0.980 | 5,970 | 22 |
| | DNABERT-2 | 0.894 | 0.896 | 0.792 | 0.950 | 0.940 | 5,819 | 22 |
| | AgroNT | **0.936** | **0.936** | **0.878** | **0.984** | **0.984** | 33,985 | 151 |
| | HyenaDNA | 0.926 | 0.918 | 0.846 | 0.968 | 0.972 | **335** | **3** |
| chromosome 3 | Motif-based | 0.744 | 0.686 | 0.524 | 0.748 | 0.728 | 3,469 | 7 |
| | DeepBind | 0.884 | 0.884 | 0.712 | 0.952 | 0.952 | 970 | 4 |
| | BERT-TFBS | 0.858 | 0.892 | 0.770 | 0.896 | 0.894 | 5,464 | 28 |
| | DNABERT-2 | 0.912 | 0.914 | 0.830 | 0.968 | 0.960 | 5,568 | 27 |
| | AgroNT | **0.965** | **0.965** | **0.928** | **0.990** | **0.990** | 34,973 | 190 |
| | HyenaDNA | 0.940 | 0.942 | 0.880 | 0.980 | 0.982 | **250** | **3.9** |
| chromosome 4 | Motif-based | 0.746 | 0.700 | 0.520 | 0.760 | 0.752 | 3,790 | 6 |
| | DeepBind | 0.882 | 0.884 | 0.768 | 0.950 | 0.950 | 1122 | 4 |
| | BERT-TFBS | 0.860 | 0.892 | 0.722 | 0.888 | 0.890 | 5,910 | 23 |
| | DNABERT-2 | 0.928 | 0.928 | 0.858 | 0.982 | 0.980 | 5,790 | 23 |
| | AgroNT | **0.966** | **0.966** | **0.934** | **0.992** | **0.994** | 36,066 | 156 |
| | HyenaDNA | 0.942 | 0.942 | 0.888 | 0.984 | 0.986 | **254** | **3** |
| chromosome 5 | Motif-based | 0.700 | 0.606 | 0.460 | 0.700 | 0.670 | 3,928 | 7 |
| | DeepBind | 0.886 | 0.886 | 0.774 | 0.952 | 0.954 | 977 | 5 |
| | BERT-TFBS | 0.952 | 0.952 | 0.904 | 0.990 | 0.990 | 5,452 | 33 |
| | DNABERT-2 | 0.918 | 0.920 | 0.840 | 0.972 | 0.972 | 5,358 | 32 |
| | AgroNT | **0.970** | **0.970** | **0.944** | **0.996** | **0.998** | 34,150 | 220 |
| | HyenaDNA | 0.944 | 0.944 | 0.884 | 0.986 | 0.984 | **251** | **4** |

#### S3.2 Cross-Dataset Evaluation on *A. thaliana* AREB/ABF2 Binding Regions

We conducted the statistical testing between HyenaDNA and the other models using Wilcoxon test.

HyenaDNA achieved the strongest performance overall, with significantly higher accuracy, F1-score, and MCC compared to all other methods based on the Wilcoxon test (all $p = 0.010$–$0.011$). For ROC-AUC, HyenaDNA significantly outperformed the Motif baseline ($p = 0.009$) and DeepBind ($p = 0.007$), while differences with BERT-TFBS, DNABERT-2, and AgroNT were not statistically significant ($p = 0.408$, $p = 0.104$, and $p = 0.498$, respectively). For PR-AUC, HyenaDNA again showed significant improvements over Motif ($p = 0.009$), DeepBind ($p = 0.009$), and DNABERT-2 ($p = 0.044$), whereas differences with BERT-TFBS and AgroNT were not significant ($p = 0.071$ and $p = 0.797$). These results underscore the superiority of HyenaDNA across most performance metrics, confirming its clear statistical advantage over both motif-based and several neural sequence models in the cross-dataset setting.

In terms of training and inference time, HyenaDNA was significantly faster than all other models, including the Motif baseline ($p = 0.008$), DeepBind ($p = 0.008$), BERT-TFBS ($p = 0.008$), DNABERT-2 ($p = 0.008$), and AgroNT ($p = 0.008$).

Table S7 summarizes the mean test-set performance metrics and runtimes (averaged over five independently trained models with different random seeds) for models trained on the `Malley2016`

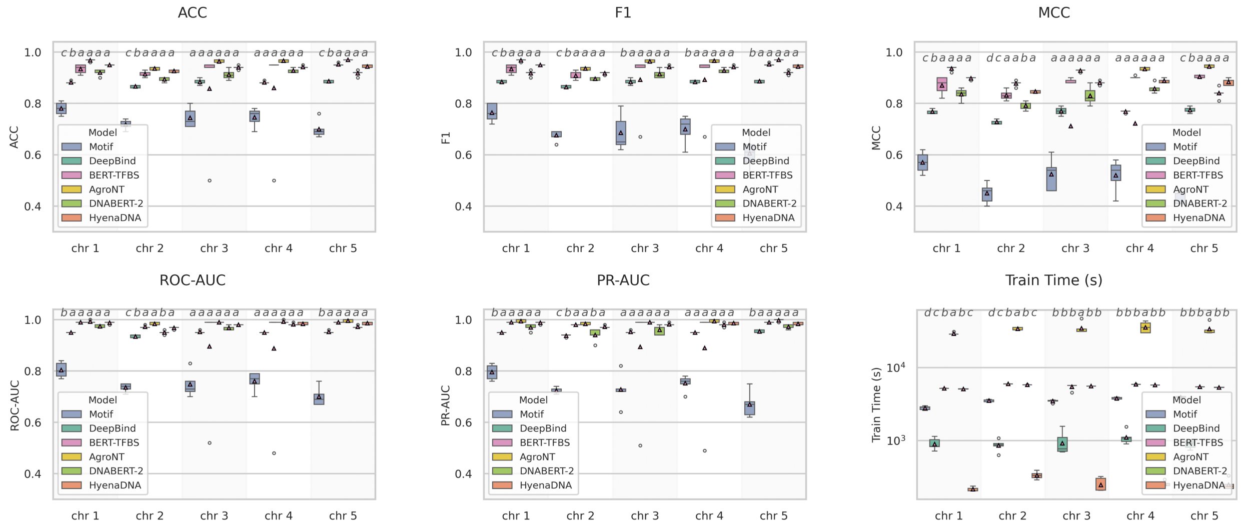


Figure S1: Per-chromosome distributions of different metrics for each model under the leave-one-chromosome-out protocol. Each marker represents the mean over five models trained with different random seeds on a single held-out chromosome. Training times are plotted on a logarithmic scale to accommodate the wide range across models. Letters on top of each box indicate statistical groupings; models sharing the same letter are not significantly different at the 0.05 significance level.

AREB/ABF2 dataset and evaluated on `Sun2022` AREB/ABF2. HyenaDNA demonstrates strong generalization, substantially outperforming classical approaches such as the motif-based method, DeepBind, and BERT-TFBS, while performing on par with DNABERT-2 and AgroNT. Notably, HyenaDNA achieves this with a fraction of the computational cost—training more than 12× faster than BERT-TFBS and DNABERT-2 and nearly 90× faster than AgroNT, and delivering inference nearly 6× faster than BERT-TFBS and DNABERT-2 and almost 40× faster than AgroNT—while maintaining competitive accuracy. These results highlight HyenaDNA as an efficient and accurate choice for cross-dataset TF-binding prediction.

Table S7: Mean performance metrics and runtimes across five independently trained models with different random seeds for the **cross-dataset** evaluation, where models were trained on *Malley2016* AREB/ABF2 and evaluated on *Sun2022* AREB/ABF2. The best value in each column is shown in bold.

| Model | ACC | F1 | MCC | ROC-AUC | PR-AUC | Train Time (s) | Test Time (s) |
|---|---|---|---|---|---|---|---|
| Motif-based | 0.786 | 0.754 | 0.590 | 0.790 | 0.774 | 3,099 | 24 |
| DeepBind | 0.804 | 0.766 | 0.638 | 0.908 | 0.920 | 162 | **16** |
| BERT-TFBS | 0.788 | 0.738 | 0.630 | 0.910 | 0.924 | 835 | 99 |
| DNABERT-2 | 0.860 | 0.844 | 0.734 | 0.908 | 0.932 | 846 | 96 |
| HyenaDNA | **0.884** | **0.876** | **0.784** | 0.942 | **0.958** | **76** | 17 |
| AgroNT | 0.840 | 0.810 | 0.716 | **0.944** | **0.958** | 6,808 | 662 |

#### S3.3 Cross-Species Transferability of AREB/ABF1-4 Binding Sites

Table S7 reports the mean test-set performance metrics and runtimes for cross-species evaluation between *A. thaliana* and *S. irio*, averaged over five independently trained models with different random seeds. HyenaDNA closely matches the predictive accuracy of leading transformer-based (e.g. DNABERT-2) and convolutional (e.g. AgroNT) baselines while training an order of magnitude faster than DNABERT-2 and over 100× faster than AgroNT. Inference time is likewise reduced by a similar factor. This combination of state-of-the-art generalization and lightweight computation makes HyenaDNA particularly well suited for rapid, large-scale transfer learning across diverse plant genomes.

Table S8: Mean performance metrics of each model for predicting AREB/ABF transcription factor binding regions in a **cross-species** setting using *A. thaliana* and *S. irio.* Metrics are averaged across five independently trained models with different random seeds. Both directions—training on one species and testing on the other—are reported. For each direction, the top value in each column is highlighted in bold.

| Direction | Model | ACC | F1 | MCC | ROC-AUC | PR-AUC | Train Time(s) | Test Time(s) |
|---|---|---|---|---|---|---|---|---|
| | Motif-based | 0.850 | 0.834 | 0.720 | 0.858 | 0.852 | 3,037 | 22 |
| **train:***A. thaliana* | DeepBind | 0.910 | 0.910 | 0.824 | 0.964 | 0.966 | 974 | 15 |
| **test:***S. irio.* | BERT-TFBS | 0.956 | 0.956 | 0.910 | **0.990** | **0.990** | 7,066 | 105 |
| | DNABERT-2 | 0.934 | 0.936 | 0.870 | 0.974 | 0.974 | 6,905 | 104 |
| | AgroNT | **0.966** | **0.966** | **0.932** | **0.990** | **0.990** | 38,344 | 717 |
| | HyenaDNA | 0.948 | 0.948 | 0.900 | 0.986 | 0.988 | **302** | **14** |
| | Motif-based | 0.708 | 0.708 | 0.426 | 0.768 | 0.754 | 3,450 | 31 |
| **train:***S.irio.* | DeepBind | 0.876 | 0.870 | 0.754 | 0.944 | 0.946 | 724 | 21 |
| **test:***A. thaliana* | BERT-TFBS | 0.926 | 0.922 | 0.852 | 0.968 | 0.978 | 5,179 | 143 |
| | DNABERT-2 | 0.918 | 0.912 | 0.830 | 0.964 | 0.964 | 5,508 | 140 |
| | AgroNT | **0.946** | **0.946** | **0.898** | **0.986** | **0.990** | 30,531 | 977 |
| | HyenaDNA | 0.928 | 0.926 | 0.854 | 0.972 | 0.974 | **213** | **20** |

### S4 Ablation of HyenaDNA Freezing Regimes

To assess how different parameter-freezing strategies affect predictive performance and computational cost of HyenaDNA, we trained the model under three regimes: freezing the entire backbone (`backbone`), freezing only the backbone layers (`backbone.layers`), and no freezing (`none`). Table S9 lists the trainable parameters under these freezing regimes:

Table S9: Trainable parameter counts for HyenaDNA under different freezing strategies.

| Frozen Components | Trainable Parameters |
|---|---|
| Backbone | 258 |
| Backbone.layers | 2,562 |
| None | 436,354 |

#### S4.1 Cross-Chromosome Evaluation

To assess the impact of parameter freezing on HyenaDNA's capacity to capture chromosome-specific features within a single species, we trained five independent models (with different random seeds) on a cross-chromosome dataset (`Sun2022` AREB/ABF transcription factor binding site samples in *A. thaliana*), where each chromosome was held out in turn for testing. Figure S2 presents the distributions average Accuracy (ACC), F1 score, Matthews correlation coefficient (MCC), ROC-AUC, and PR-AUC across five leave-one-chromosome-out experiments for three finetuning strategies. Models trained without freezing (pink) achieve the best performance across all metrics, demonstrating strong within-species generalization. Freezing only the backbone layers (green) results in a modest reduction, suggesting that key representations are retained while still allowing adaptation. In contrast, freezing the entire backbone (purple) substantially impairs performance, highlighting the importance of updating at least part of the pretrained parameters even in within-species settings.

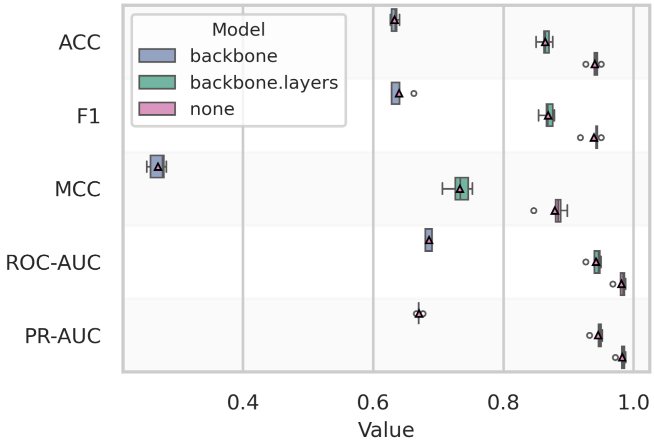


Figure S2: Distribution of test-set performance metrics for three parameter-freezing strategies in the HyenaDNA model, assessed using **five cross-chromosome hold-out** runs. For each cross-chromosome hold-out, five models trained with different random seeds and test set were evaluated. Boxplots show the distribution of mean Accuracy (ACC), F1 score, Matthews correlation coefficient (MCC), area under the ROC curve (ROC-AUC), and area under the precision–recall curve (PR-AUC) across the cross-chromosome hold-outs. Purple, green, and pink represent freezing the entire backbone, freezing only the backbone layers, and no freezing, respectively. Mean results for individual runs across the five runs are shown as circles, while averages are indicated with triangles.

#### S4.2 Cross-Dataset Evaluation

We trained the HyenaDNA model under the three regimes using cross-dataset experiment in which models trained on the `Malley2016` AREB/ABF2 dataset were tested on the independent `Sun2022` *A. thaliana* AREB/ABF2 dataset. We compared three finetuning regimes: (i) freezing the entire pretrained backbone, (ii) freezing only the backbone layers, and (iii) full finetuning (no freezing). Performance was measured over five independent models (with different random seeds), reporting Accuracy (ACC), F1 score, Matthews correlation coefficient (MCC), ROC-AUC and PR-AUC for each run (Figure S3).

When all parameters were trainable (pink), HyenaDNA adapted most effectively to the new dataset, achieving the highest mean scores across all metrics. Partial freezing of backbone layers (green) yielded intermediate performance, indicating some preservation of pretrained features but reduced flexibility to dataset-specific signal. Fully freezing the backbone (purple) severely degraded cross-dataset generalization, specially in terms of MCC, underscoring that without any parameter updates HyenaDNA cannot accommodate differences between protocols. These results demonstrate that model finetuning is essential for high-fidelity prediction when transferring between distinct experimental datasets.

#### S4.3 Cross-Species Evaluation

We performed two leave-one-species-out experiments on *A. thaliana* and *S.irio* species using `Sun2022` AREB/ABF1-4 peak regions under the three regimes (Figure S4). In each case, five different seeds were used for training the model on one species and testing on the other. When all parameters were unfrozen (pink), the model attained excellent transfer performance regardless of training direction. By contrast, freezing the entire backbone (purple) almost completely abrogated cross-species prediction, showing that without any parameter updates the model cannot adapt to species-specific

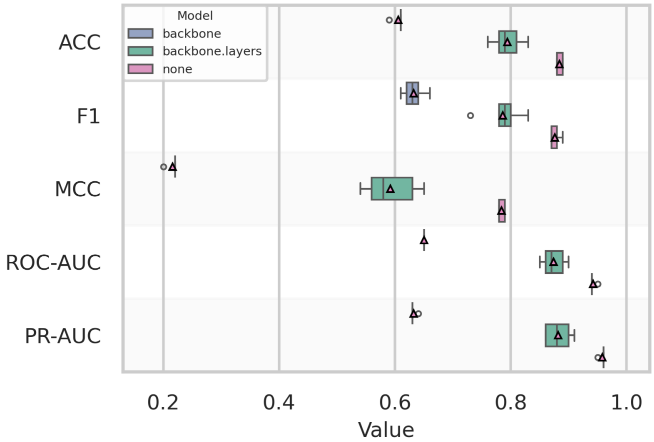


Figure S3: Distribution of test-set performance (ACC, F1, MCC, ROC-AUC, PR-AUC) per trained model for finetuning strategies of HyenaDNA in the **cross-dataset** evaluation (models trained on `Malley2016` AREB/ABF2, tested on `Sun2022` *A. thaliana* AREB/ABF2). Purple, green and pink correspond to freezing the entire backbone, freezing only backbone layers, and no freezing, respectively. Individual results are overlaid as circles and means as triangles

signals. Selectively freezing only the backbone layers (green) provided an intermediate solution: it recovered most of the pretrained knowledge while still allowing adaptation. These results underscore that end-to-end finetuning is essential for robust cross-species generalization, and that partial backbone freezing offers a favourable trade-off between leveraging pretrained representations and adapting to new genomic contexts.

(a)

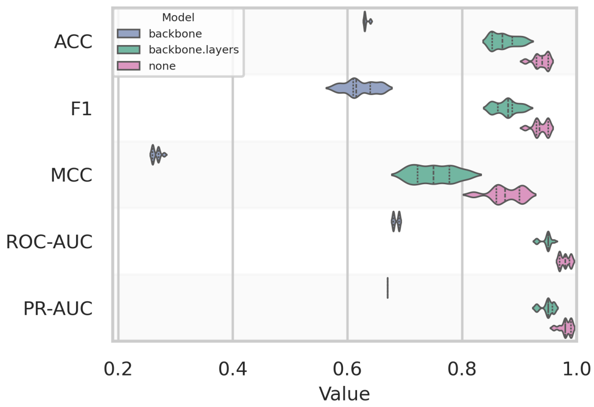


(b)

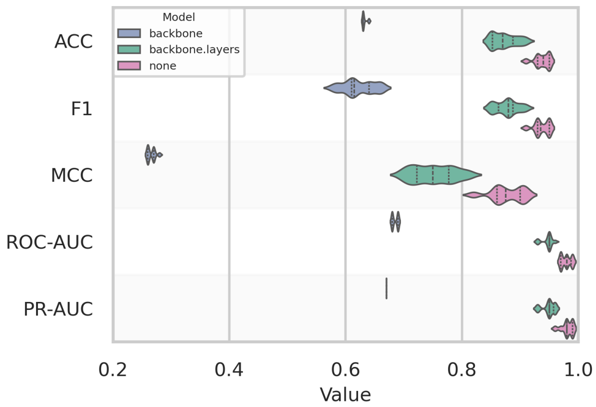


Figure S4: Distribution of test-set performance metrics for three parameter-freezing strategies in the HyenaDNA model under **cross-species** evaluation. Each distribution summarizes results from five independently trained models (with different random seeds), where models are evaluated on the held-out test species. **(a)**: trained on *A. thaliana*, tested on *S. irio*; **(b)**: trained on *S. irio*, tested on *A. thaliana*. Purple, green and pink correspond to freezing the entire backbone, freezing only backbone layers, and no freezing, respectively. Individual results are overlaid as circles and means as triangles.

[7] Ilya Loshchilov and Frank Hutter. Decoupled weight decay regularization. arXiv preprint arXiv:1711.05101, 2017.
